# Localized hypermutation drives the evolution of unstable colistin resistance in *Pseudomonas aeruginosa*

**DOI:** 10.1101/2021.08.10.455869

**Authors:** Natalia Kapel, Julio Diaz Caballero, R. Craig MacLean

## Abstract

Colistin has emerged as an important last line of defence for the treatment of infections caused by antibiotic resistant Gram-negative pathogens. Here we investigate the responses of ≈1,000 populations of an MDR strain of *P. aeruginosa* to a high dose of colistin. Colistin exposure resulted in rapid cell death, but a sub-set of populations eventually recovered due to the outgrowth of heteroresistant cells. Genome sequencing revealed that heteroresistance was primarily driven by mutations in the PmrAB two-component system that occurred at a rate (≈2×10^-5^ per cell division) that was 10^3^-10^4^ fold higher than typical resistance mutation rates. Crucially, this elevated mutation rate was only found in *pmrB*, demonstrating that hypermutability is localized to this gene. PmrAB provides resistance to antimicrobial peptides that are involved in host immunity, suggesting that this pathogen may have evolved a high mutation rate as an adaption to generate mutants that are resistant to host antimicrobial peptides that are secreted during infection. Interestingly, we found no mutations in 1/3 of populations that recovered from colistin treatment, suggesting that phenotypic plasticity and/or persister cells contribute to the ability of *Pseudomonas* to adapt to colistin.

## Introduction

The antimicrobial peptide colistin has emerged as an important last line of defense for the treatment of infections caused by multi-drug resistant (MDR) Gram-negative pathogens, including *Escherichia coli, Klebsiella pneumoniae, Pseudomonas aeruginosa* and *Acinetobacter baumanii*[1–3]. Given this, there is a pressing need to understand how bacterial pathogens adapt to colistin treatment.

Unfortunately, colistin has moderate efficacy for the treatment of *P. aeruginosa* infections, in spite of the fact that resistance, as defined by clinical breakpoint MICs, remains rare[3–6]. One possible explanation for this is that poor pharmacodynamics prevent colistin from achieving high enough concentrations at the sites of *P. aeruginosa* infection[2, 7]. It is also possible that bacterial responses to colistin contribute to treatment failure. For example, treatment failure could be driven by adaptive changes in gene expression that confer colistin resistance *in vivo*[8], by the growth of persister cells that survive colistin treatment[9], or by heteroresister cells that carry unstable resistance mutations[10]. Importantly, in all these cases we would not expect to see any stable increase in the resistance of isolates recovered after antibiotic treatment, as standardized methods for resistance testing rely on pre-culturing bacterial isolates in antibiotic-free medium.

The classical model for the evolution of resistance during infection is that antibiotic treatment selects for rare *de novo* resistance mutations that confer a stable antibiotic-resistant phenotype that is associated with fitness costs[12–14]. Given that resistance mutations occur at a low rate, this model is most applicable to cases where infection results in a large bacterial population, or when bacterial mutation rates are high. Large populations of *P. aeruginosa* adapt to high concentrations of colistin in the lab through the sequential fixation of a series of mutations, generating elevated resistance[15, 16]. Whilst this approach offers a powerful and elegant tool for understanding evolutionary trajectories and constraints, the clinical relevance of these experiments is unclear given that colistin resistance is rare in clinical settings.

Our approach to study responses to colistin was to expose ≈1,000 small populations (10^4^-10^5^ cells) of *P. aeruginosa* EP67, a colistin-sensitive MDR isolate[11] to the clinical breakpoint dose of colistin (2mg/L). In this selective regime, populations could only persist over the long term if they were able to acquire increased antibiotic resistance. Our reasoning for this approach was that using many small populations of bacteria prevents rare mutations (i.e. 10^-8^-10^-9^ per cell division[17, 18]) that confer high level resistance from dominating the evolutionary response to antibiotic treatment[19]. A further advantage of this approach was that studying responses to antibiotic treatment in small populations opened up the possibility that bacterial populations were driven to extinction before resistance could evolve[20, 21], which is the most common outcome of antibiotic treatment in the clinic[3, 5, 6, 22].

## Material and Methods

### Strain and culture conditions

In this study we used strain EP67, an MDR strain of *P. aeruginosa* ST17 that was isolated from an ICU patient with ventilator-associated pneumonia[11]. The patient that this strain was isolated was treated with colistin, but this isolate was collected prior to colistin treatment. Unless otherwise stated, cultures were grown at 37°C in Luria-Bertani (LB) medium with shaking at 250 rpm or, alternatively, statically and solidified with 1% agar when appropriate. For all assays, colistin was freshly prepared from 5mg/mL powder in water solution.

### Colistin time-kill experiment

Glycerol stocks of Strain EP67 were streaked out on LB plates and single isolated colonies were inoculated into 200μL of LB Miller and grown over 18-20h at 37°C with shaking at 225rpm. Overnight cultures were diluted in PBS and inoculated into 96 well polystyrene microtiter plates containing 200μL of fresh LB Miller containing colistin at a concentration of 2mg/L, such that the initial cell density was ≈1×10^6^ CFU/mL (i.e. 2×10^5^CFU/culture). Colistin cultures were then incubated at 37°C with shaking at 225 rpm. Samples of colistin cultures were serially diluted and spotted (5μL) on LB agar plates and CFUs were counted after overnight incubation at 37°C. The minimal dilution factor that was used was 10-fold, to ensure that colistin that was carried over to LB plates was at a sub-MIC dose (i.e. <0.2mg/L). 100 independent replicates of this experiment were carried out across 4 different blocks. Viable cell density at times 0-8 hr was estimated from the proportion of cultures that gave 0 CFUs at a dilution factor of 0.001 (time=0) or 0.1 (all other time points) under the assumption that the number of CFUs/sample follows a Poisson distribution. Confidence intervals in the proportion of CFU-free samples were estimated from the normal approximation to the binomial distribution.

### Large-scale rebounding experiment

Cultures of strain EP67 were pre-cultured and challenged with colistin using the same methods as in the time-kill experiments above, except that a lower titre of cells was inoculated into colistin containing medium (≈5×10^4^ CFU/mL (i.e. 1×10^4^CFU/culture)). This experiment was replicated 933 times across 6 experimental blocks. Initial cell titre in each block was determined by serially diluting and plating samples from 60 independent cultures on LB and counting CFUs after overnight incubation. All cultures were diluted and spotted onto LB plates after 24 and 48 hrs of incubation with colistin to quantify the frequency of population rebounding.

### Testing for heteroresistance

In this experiment, we challenged 171 independent cultures of EP67 with 2mg/L colistin, as in the time-kill experiment. After 24 hrs of incubation in colistin, each selection culture was diluted 10-fold into fresh LB medium, and these ‘recovery cultures’ were incubated overnight. Note that a 10-fold dilution was sufficient to reduce the concentration of colistin to a sub-MIC level (0.2mg/L) that does not select for colistin resistance. After overnight incubation, recovery cultures were then passaged into (i)LB and (ii)LB+colistin (2mg/L) by a 1000-fold serial dilution to set up ‘test cultures’ that were incubated overnight. After overnight incubation, all of the ‘recovery cultures’ and ‘test cultures’ were diluted 10-fold and 5 μL dots were plated on LB plates that were incubated overnight. Populations were counted as viable if they produced confluent colony growth, which corresponds to a culture density of at least 16,000 CFUs.

### DNA Extraction

DNA was extracted from 18 cultures that were growing in LB+colistin (2mg/L) using DNeasy Blood and Tissue Kit (QIAGEN)

### Sequencing and Bioinformatic analysis

All samples were sequenced in the MiSeq illumina platform using the 300×2 Paired End protocol. The sequencing experiment yielded a coverage of 64X-168X. Raw reads were quality controlled with Trimmomatic v. 0.39[23] using the ILLUMINACLIP (2:30:10) and SLIDING WINDOW (4:15) parameters. The coverage of the resulting quality-controlled reads ranged from 51X to 131X. Small variants were identified using breseq v. 0.34.0[24] in the polymorphism prediction mode, which implemented bowtie2 v. 2.3.5.1[25]. Copy number variants were estimated with the default parameters of the package CNOGpro v. 1.1[26] in *R*[27] using the *P. aeruginosa* reference genome EP67 (SAMN16363815, https://www.ncbi.nlm.nih.gov/biosample/16363815).

### Mutation rate estimation

The mutation rate at *pmrB* was estimated using the method of Luria and Delbrück[28]. We estimated the proportion of cultures containing *pmrB* mutants by multiplying the number of cultures that tested positive for growth after 48hrs of incubation in colistin by the proportion of sequenced cultures containing *pmrB* mutations (8/15). The mutation rate at *pmrB* was estimated as -lnp(0)/N, where p(0) is the estimated proportion of cultures that did not contain a *pmrB* mutant and *N* is the initial number of CFU in the culture. The mutation rate was estimated for the initial time kill experiment and for the 6 blocks of the large scale rebound experiment. The error in the mutation rate was estimated as the standard error of the 7 independent estimates of the mutation rate.

## Results

### PA responses to colistin treatment

To investigate the response of *Pseudomonas* populations to colistin therapy, we inoculated 100 independent replicate cultures of EP67 into fresh culture medium containing the clinical breakpoint concentration of colistin (2mg/L). Viable cell density rapidly declined and after 8 hours of incubation we recovered viable colonies from only 2/100 cultures, giving an estimated cell density of only 10 CFUs/culture. The number of cultures that gave detectable growth increased to 15%-20% after continued incubation in colistin containing medium, demonstrating that a small subset of EP67 populations eventually recovered from colistin treatment. The viable cell density in populations showing evidence of recovery increased between 24 hours and 48 hours (rank-sum test, Z=1.99, P=0.046), but remained very low, with densities typically <5000 CFU (Figure 1C). Interestingly, this low cell density is well below the number of cells required to form visible colonies (>10^6^CFU) that are usually used to detect the growth of antibiotic resistant populations.

**Figure 1:**
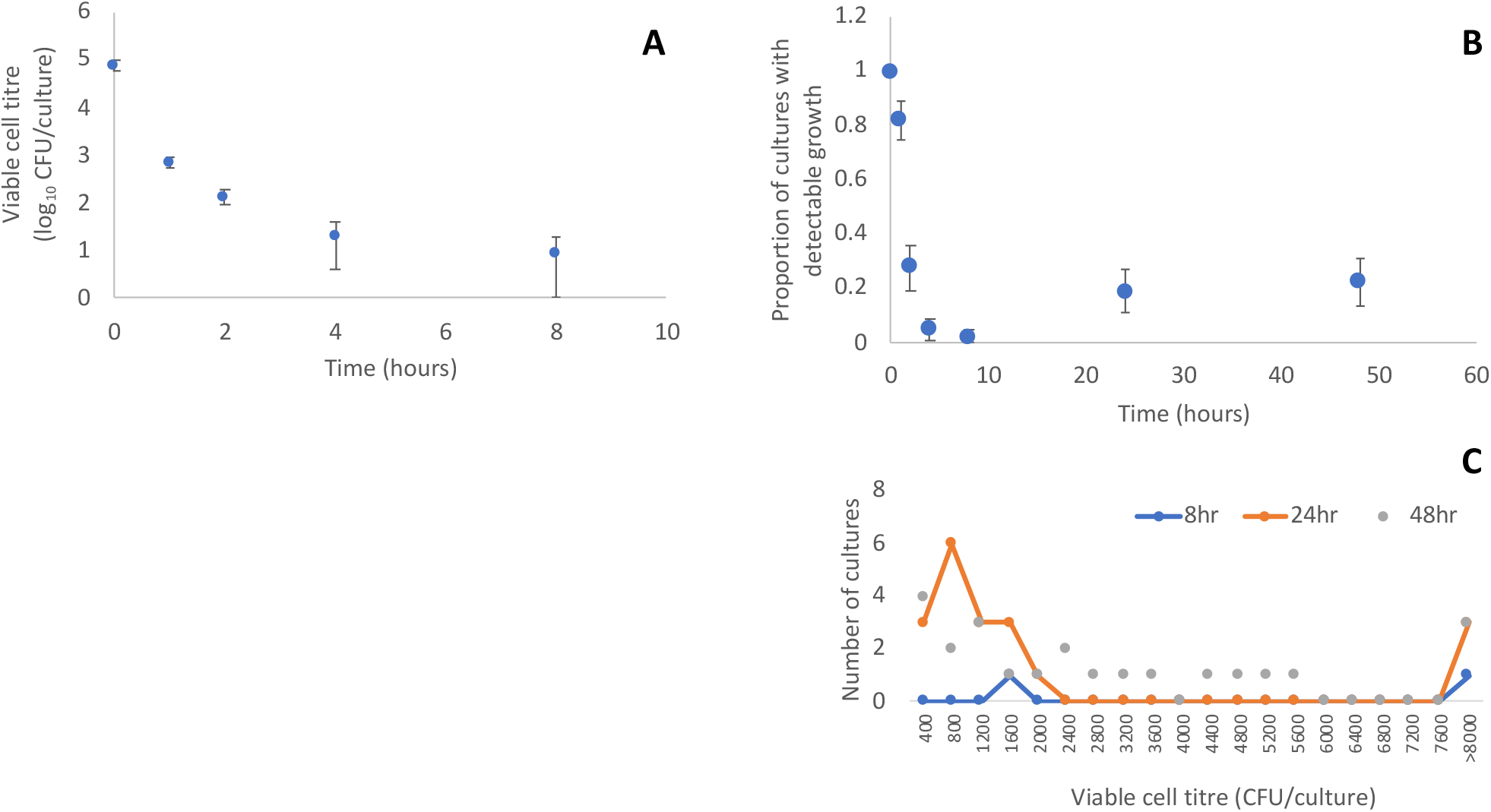
Population responses to colistin treatment. 100 independent populations of strain EP67 were inoculated into culture medium supplemented with colistin (2mg/L). (**A**) Viable cell titre (+/- 95% c.i) over the first 8 hours of incubation. (**B**) Proportion of cultures (+/- 95% c.i) showing detectable growth over time, with a minimal limit of detection of 400 CFU/culture. (**C**) Distribution of viable cell titre in cultures showing growth at 8hr, 24hr and 48 hr. The limits of detection in this assay were 400-8000 CFUs.

The low frequency of recovery of PA populations after colistin treatment suggests that populations recovery was driven by rare events, such as spontaneous mutation or gene amplification. To better understand the frequency and repeatability of population recovery, we repeated this assay 6 times, with a total of 933 independent cultures. In these large-scale assays, we focused only on measuring the density of viable cells after 24 and 48hrs of incubation. Population recovery was detected in all assays, but the frequency of recovery differed quite widely between assays (Table 1).

**Table 1:** Summary of population rebounding experiments. This table shows a summary of the frequency of *Pseudomonas* population recovery in replicate colistin challenge experiments.

| Experiment | Number of replicate cultures | Initial CFU/culture | Number of viable cultures |  |
| --- | --- | --- | --- | --- |
|  |  |  | 24 hrs | 48 hrs |
| Time kill | 100 | 7.09E+04 | 19 | 23 |
| Rebounding 1 | 171 | 5.62E+03 | 6 | 9 |
| Rebounding 2 | 162 | 1.67E+04 | 64 | 52 |
| Rebounding 3 | 60 | 7.14E+03 | 2 | 32 |
| Rebounding 4 | 180 | 6.71E+03 | 54 | 99 |
| Rebounding 5 | 180 | 9.20E+03 | 37 | 75 |
| Rebounding 6 | 270 | 8.68E+03 | 13 | 17 |

Population recovery in the presence of colistin could have been driven by the growth of either classical resistant mutants[15, 16] or heteroresistant cells[29]. To discriminate between these possibilities, we carried out a further experiment to measure the stability of colistin resistance (Figure 2). In this experiment, we first selected for colistin resistance, as in our previous experiments. Populations that were selected for colistin resistance were then passaged into colistin-free culture medium to allow populations that survived treatment to expand. These recovered populations were then passaged into medium containing colistin, to measure the stability of the colistin resistant phenotype, and colistin-free medium, as a control. Colistin resistance was only maintained in 2 of the 56 populations (3.5%) that recovered from initial colistin treatment, allowing us to rule out the possibility that population recovery was driven by classical resistance mutations.

**Figure 2:**
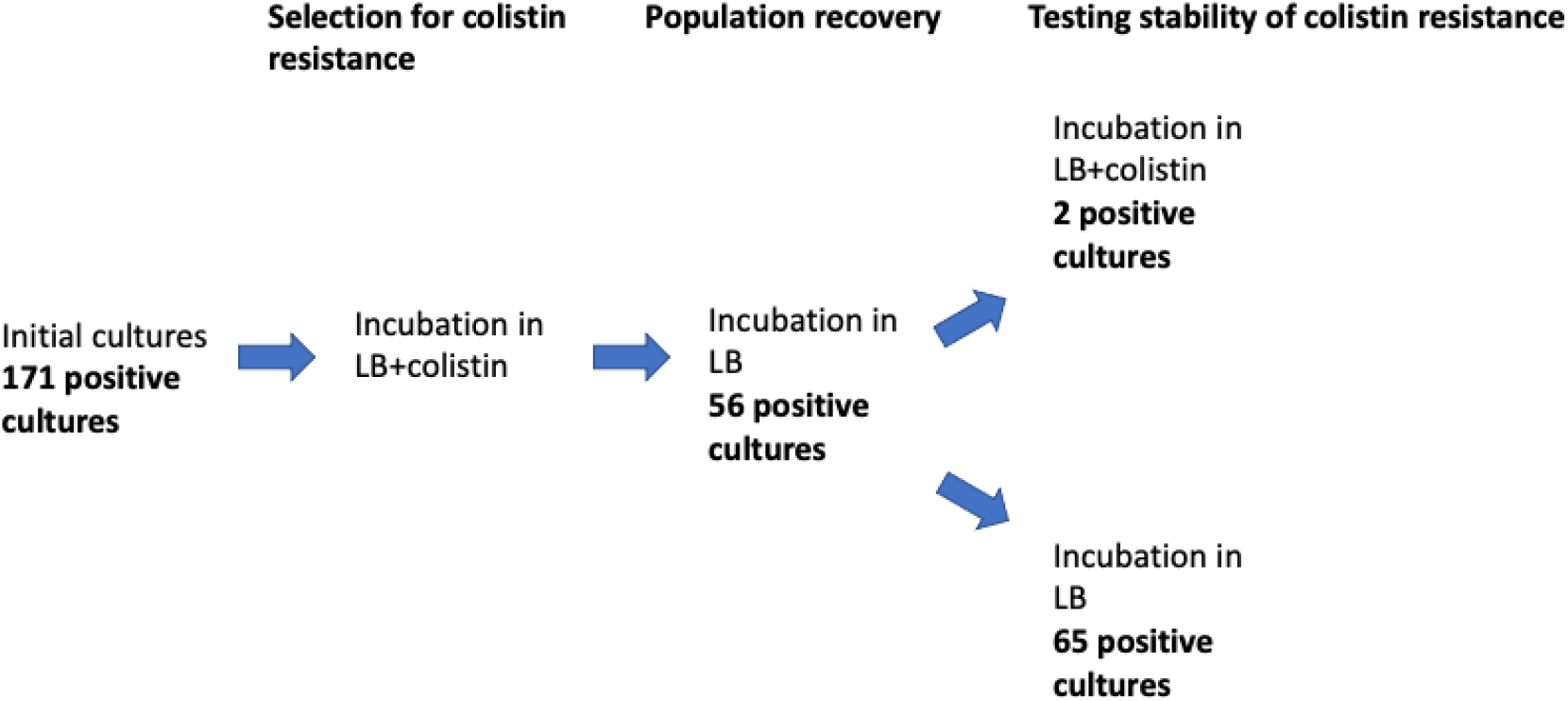
Testing the stability of colistin resistance. 171 cultures of EP67 were incubated with colistin for 24 hours, to select for resistance. Cultures were then transferred to LB, to allow resistant populations to expand. Recovered populations were then passaged into medium with LB or LB+colistin to test the stability of colistin resistance. After each passage, cultures diluted 10 fold and spotted out on LB agar plates to determine population viability, as determined by confluent colony growth.

### Hypermutability of *pmrB* drives heteroresistance

Heteroresistance is driven by unstable genetic changes (i.e. gene amplification, and SNPs) where both forward and reverse mutation occur at a high rate, ensuring the existence of high levels of polymorphism in a localized region of the genome[10]. A classic example of this phenomenon is when a gene that confers resistance undergoes rapid changes in copy number due to the presence of repeated flanking regions, such as transposons[18]. Given that antibiotic resistance tends to be costly, an important consequence of heteroresistance is that resistance is lost from bacterial populations following antibiotics treatment, as we observed.

To directly test for HR, we sequenced 18 cultures that were growing in medium containing colistin (Table 2). Sequencing revealed that 3/18 cultures were *Micrococcus luteus*, implying that some contamination occurred during our experiments. *M. luteus* contamination was associated with a very conspicuous phenotype (yellow colonies) that was only detected very sporadically, allowing us to rule out the possibility that *M. luteus* contamination was common. The remaining 15 sequenced cultures contained SNPs at two hot-spot sites in *pmrB* (n=8), which is part of a two-component regulatory system that has been previously implicated in colistin heteroresistance[30–32], or short amplifications of genes involved in LPS biosynthesis (n=2). Remarkably 5/15 cultures matched perfectly to the EP67 reference genome and did not contain any SNPs or gene amplifications.

**Table 2:**
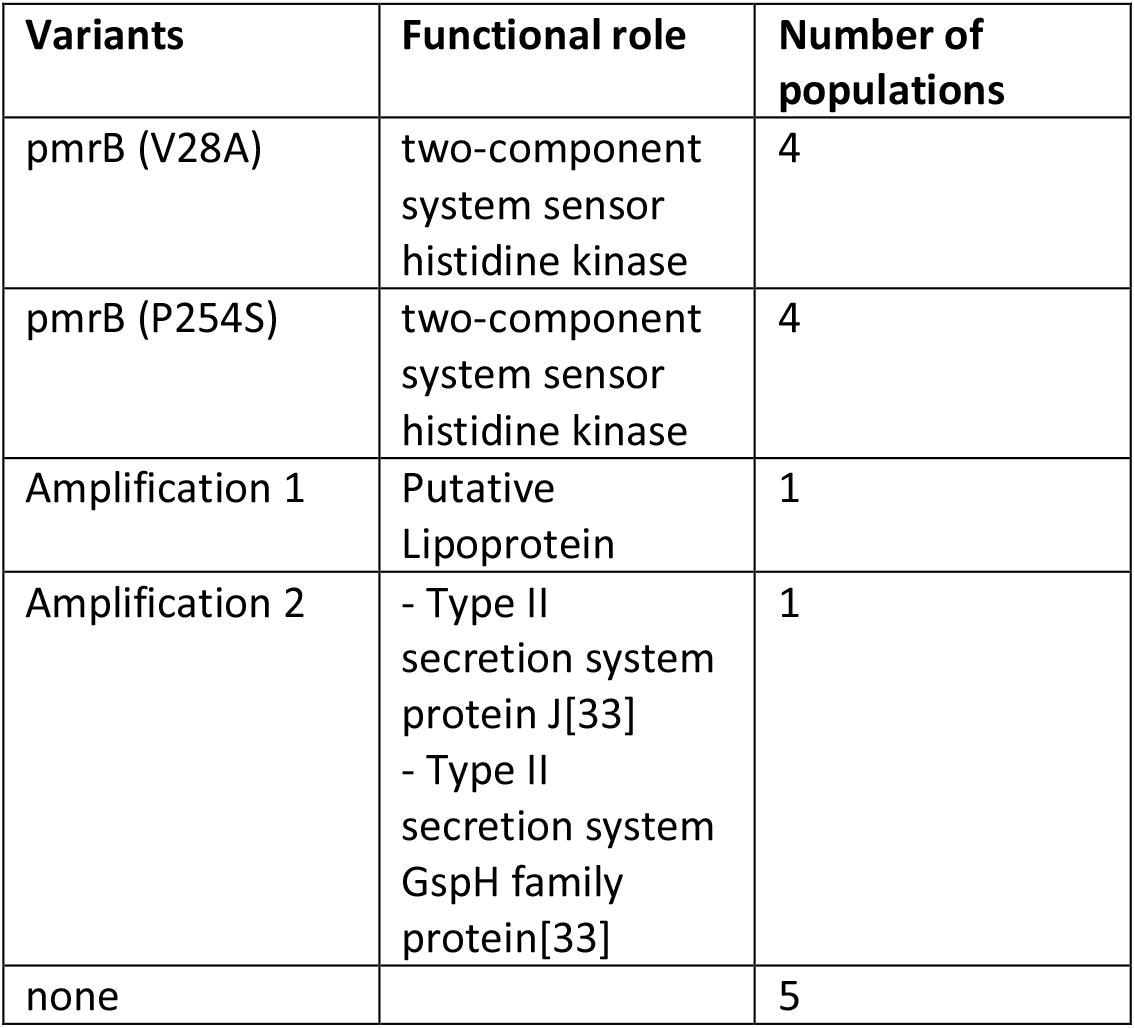
Summary of detected variants in rebounding populations.

Given that HR is driven by mutation, it is possible to use existing methods to estimate the underlying rate at which heteroresistance occurs. We used the Luria-Delbrück[28, 34] method to estimate the rate of occurrence of HR mutations. This simple method, which assumes that mutants are already present at the time of selection (in this case, treatment with colistin), estimates the mutation rate based on the cell titre and the frequency of cultures lacking mutants of interest. In this case, we assumed that the frequency of *pmrB* mutants was 8/15 across all of the 7 rebounding experiments (Table 1). Given this assumption, we estimate that *pmrB* mutations occurred 2×10^-5^ per cell division (s.e.=7.96×10^-6^; n=7 assays). Note that this is a conservative estimate, as our mutation rate estimates are based on the proportion of populations with viable cells after 48hrs of incubation. Because we only sampled a fraction of each population at this time point, this must underestimate the true frequency of viable populations.

## Conclusion

In this study, we explored evolutionary responses to colistin treatment in many small populations of *P. aeruginosa*. Colistin treatment led to rapid cell death, and population size declined to approximately 10 cells after 8 hours of colistin treatment (Figure 1). Remarkably, approximately 30% of populations recovered (Figure 1, Table 1) due to the slow growth of heteroresister cells with an unstable colistin resistance phenotype (Figure 2). The dominant genetic mechanism of HR was mutations in PmrAB (Table 2), a two-component regulatory system that modifies LPS through the addition of aminoarabinose to Lipid A[32]. HR was also driven by small amplifications of genes involved in LPS biosynthesis, providing further evidence for the importance of LPS alterations to colistin resistance. There are many known examples of heteroresistance to colistin[10, 30, 31, 36–38], and the PmrAB system has been implicated in colistin resistance in many studies[15, 16, 30–32, 39]. The key insight from our study is that the hypermutability of *pmrB* allows small *P. aeruginosa* populations to adapt to colistin treatment.

Stressful conditions, including antibiotics, are known to increase the mutation rate in bacteria[40]. Given this, is possible that the mutagenic effects of colistin lead to mutations in *pmrB*. However, stress induced mutagenesis creates mutations across the genome[41], and the absence of second-site mutations provides good evidence that *pmrB* mutants were present at the time of colistin exposure due to the fact that this gene has an inherently high mutation rate. The mutation rate of *pmrB* is extraordinarily high (at least 2×10^-5^per cell division), given that bacterial mutation rates are typically on the order of 10^-10^ per base pair per division[42], and chromosomal resistance mutations typically arise at a rate of 10^-8^-10^-9^ per cell division[17, 18].

Interestingly, mutations in the PmrAB system have been shown to facilitate airway colonization by *P. aeruginosa* and provide resistance to host antimicrobials that are produced as part of the innate immune responses, such as LL37 and lysozyme[32, 43]. *P. aeruginosa* is an opportunistic pathogen, and our results raise the intriguing possibility that PmrAB may have evolved a high mutation rate to allow rapid adaptation to the host environment[44] in a way that is analogous to the contingency loci that generate phase variation in *Haemophilus* and *Neisseria*[45, 46]. Given the instability of the *pmrB* colistin resistance phenotype, a key goal for future work will be to determine the extent to which *pmrB*-mediated HR drives treatment failure in patients undergoing colistin therapy for *P. aeruginosa* infections. Although *pmrB* mutations provide a simple mechanism for *P. aeruginosa* to adapt to colistin, *pmrB* mutations are known to have pleiotropic effects, including enhanced antibiotic susceptibility[43], that may effectively select against *pmrB in vivo*.

One of our most surprising findings was that no mutations were identified in 5/15 populations that recovered under colistin treatment. Given the depth of coverage of Illumina sequencing in these populations and the high quality of the EP67 reference genome that was used for read-mapping, we can be confident that these represent true negatives and not false negative where we were unable to identify mutations. One possible explanation is that mutation-free populations reflects the growth of persister cells[9]. It is challenging to test this hypothesis, but it is difficult to see how populations of persister cells could continue to grow under colistin treatment (Figure 1C). An alternative explanation is that phenotypic plasticity contributes to colistin heteroresistance. According to this explanation a small sub-set of *P. aeruginosa* cells are able to actively grow in the presence of colistin due to either inherent or induced[8] expression of genes that modify the membrane to make it resistant to antimicrobial peptides like colistin. An important goal for future work will be to rigorously test the non-genetic mechanisms that allow *P. aeruginosa* populations to recover under colistin treatment.

## Statement of author contributions

The project was conceived by NK and RCM. NK carried out experimental work. All authors contributed to data analysis and interpretation. RCM wrote the manuscript, which was edited by NK and JDC.

## Acknowledgements

This research was supported by Wellcome Trust Grant (106918/Z/15/Z). We thank the Oxford Genomics Centre (funded by Wellcome Trust Grant 203141/Z/16/Z) for the generation and initial processing of Illumina sequence data.

